# SARS-CoV-2 variants of concern have acquired mutations associated with an increased spike cleavage

**DOI:** 10.1101/2021.08.05.455290

**Authors:** Alba Escalera, Ana S. Gonzalez-Reiche, Sadaf Aslam, Ignacio Mena, Rebecca L. Pearl, Manon Laporte, Andrea Fossati, Raveen Rathnasinghe, Hala Alshammary, Adriana van de Guchte, Mehdi Bouhaddou, Thomas Kehrer, Lorena Zuliani-Alvarez, David A. Meekins, Velmurugan Balaraman, Chester McDowell, Jürgen A. Richt, Goran Bajic, Emilia Mia Sordillo, Nevan Krogan, Viviana Simon, Randy A. Albrecht, Harm van Bakel, Adolfo Garcia-Sastre, Teresa Aydillo

**Affiliations:** Department of Microbiology, Icahn School of Medicine at Mount Sinai; New York, NY, USA; Global Health and Emerging Pathogens Institute, Icahn School of Medicine at Mount Sinai; New York, NY, USA; Graduate School of Biomedical Sciences, Icahn School of Medicine at Mount Sinai; New York, NY, USA; Department of Genetics and Genomic Sciences, Icahn School of Medicine at Mount Sinai; New York, NY, USA; Quantitative Biosciences Institute (QBI), University of California San Francisco; San Francisco, CA, USA; QBI COVID-19 Research Group (QCRG); San Francisco, CA, USA; J. David Gladstone Institutes; San Francisco, CA, USA; Department of Cellular and Molecular Pharmacology, University of California San Francisco; San Francisco, CA, USA; Department of Diagnostic Medicine/Pathobiology and Center of Excellence for Emerging and Zoonotic Animal Diseases, College of Veterinary Medicine; Kansas State University, Manhattan, KS, USA; Department of Pathology, Molecular and Cell-Based Medicine, Icahn School of Medicine at Mount Sinai; New York, NY, USA; Department of Medicine, Division of Infectious Diseases, Icahn School of Medicine at Mount Sinai; New York, NY, USA; Icahn Institute for Data Science and Genomic Technology, Icahn School of Medicine at Mount Sinai; New York, NY, USA; The Tisch Cancer Institute, Icahn School of Medicine at Mount Sinai; New York, NY, USA

## Abstract

For efficient cell entry and membrane fusion, SARS-CoV-2 spike (S) protein needs to be cleaved at two different sites, S1/S2 and S2’ by different cellular proteases such as furin and TMPRSS2. Polymorphisms in the S protein can affect cleavage, viral transmission, and pathogenesis. Here, we investigated the role of arising S polymorphisms *in vitro* and *in vivo* to understand the emergence of SARS-CoV-2 variants. First, we showed that the S:655Y is selected after *in vivo* replication in the mink model. This mutation is present in the Gamma Variant Of Concern (VOC) but it also occurred sporadically in early SARS-CoV-2 human isolates. To better understand the impact of this polymorphism, we analyzed the *in vitro* properties of a panel of SARS-CoV-2 isolates containing S:655Y in different lineage backgrounds. Results demonstrated that this mutation enhances viral replication and spike protein cleavage. Viral competition experiments using hamsters infected with WA1 and WA1-655Y isolates showed that the variant with 655Y became dominant in both direct infected and direct contact animals. Finally, we investigated the cleavage efficiency and fusogenic properties of the spike protein of selected VOCs containing different mutations in their spike proteins. Results showed that all VOCs have evolved to acquire an increased spike cleavage and fusogenic capacity despite having different sets of mutations in the S protein. Our study demonstrates that the S:655Y is an important adaptative mutation that increases viral cell entry, transmission, and host susceptibility. Moreover, SARS-COV-2 VOCs showed a convergent evolution that promotes the S protein processing.

## Introduction

SARS-CoV-2 has been spreading worldwide causing millions of infections and deaths since its emergence in Wuhan, China, in late 2019. Apart from humans, ferrets, cats, dogs, Syrian golden hamsters, and nonhuman primates are also susceptible to SARS-CoV-2 infection and transmission (*1, 2*). In addition, cases of viral spread in mink farms and mink-to-human cross-species transmission have been reported (*3, 4*). The spike (S) glycoprotein of SARS-CoV-2 is the main determinant of host tropism and susceptibility, and the main target of antibody responses (*5*). Therefore, the emergence of adaptive mutations present in the spike protein can strongly affect host tropism and viral transmission (*6, 7*). The S protein is composed of two subunits: S1 which contains the receptor binding domain (RBD) that initiates infection by binding to the angiotensin converting enzyme 2 (ACE2) receptor present in the host cell surface; and the S2 subunit that mediates fusion between viral and cellular membranes (*8, 9*). To fuse with the host cell, the S protein needs to be cleaved by cellular proteases at the S1/S2 and S2’ sites. Importantly, the S1/S2 site of SARS-CoV-2 viruses contains a multibasic furin motif (_681_PRRXR_685_) absent in other beta coronaviruses (*10, 11*) that can be processed by furin proteases, but also by transmembrane serin proteases such as TMPRSS2, or by cathepsins present in the endosomes (*10, 12-15*). The S1/S2 cleavage exposes the S2’ site, and a second cleavage of the S2’ is needed to release an internal fusion peptide that mediates membrane fusion (*16*).

Since 2019, several SARS-CoV-2 lineages have emerged leading to the divergence of an extensive subset of more transmissible SARS-CoV-2 variants termed Variants of Concern (VOCs). This has led to the natural selection of several mutations in the spike protein with different functional consequences, some of them unknown. As SARS-CoV-2 variants are raising, more research is needed to understand what the drivers of evolutionary changes are over time, and the potential impact on the epidemiology, antigenicity, escape from neutralizing antibodies induced by previous infection and vaccination and virus fitness. The first widely adaptative substitution described was the S: D614G which became dominant in March 2020 and is present in most of the variants currently circulating worldwide. This substitution is known to enhance viral replication in the upper respiratory tract as well as *in vivo* transmission (*17, 18*). Several other polymorphisms became dominant in late 2020. The N501Y substitution convergently evolved in early emerging VOCs Alpha (B.1.1.7), Beta (B.1.351) and Gamma (P.1) variants and has been associated with an enhanced spike affinity for the cellular ACE2 receptor (*19, 20*). This mutation is located in the receptor-binding motif (RBM) of the RBD, the primary target of many neutralizing antibodies. Importantly, accumulation of mutations in the RBD can decrease neutralizing antibody responses elicited by infection or vaccination against ancestral SARS-CoV-2 variants (*21*). Similarly, the later SARS-CoV-2 Kappa (B.1.617.1) and Delta (B.1.617.2) variants have also shown a significantly reduced sensitivity to convalescent and immune sera (*22-24*). Other mutations outside the RBD have also become prevalent. A clear example is the polymorphism found at position S:681 in the furin cleavage site, which includes P681H and P681R in the Alpha and Kappa/Delta variants, respectively. Some preliminary reports have pointed to an enhancement in virus transmissibility associated with this polymorphism, perhaps due to an increase of spike cleavage (*25*). Additionally, several other mutations have been identified at the edge of the furin cleavage site. This is the case of the H655Y substitution found in the Gamma (P.1) variant. This mutation was associated with changes in antigenicity by conferring escape from human monoclonal antibodies (*26*). Moreover, it has also been found to be selected in animal models after experimental infection *in vivo* (*27, 28*), indicating a potential role in host replication, transmissibility, and pathogenicity.

Here, we characterized emerging SARS-CoV-2 spike polymorphisms *in vitro* and *in vivo* to understand their impact on transmissibility, virus pathogenicity and fitness. Using the mink model of COVID-19, we found that the S:H655Y substitution was acquired *in vivo* after infection with the WA1 isolate (USA-WA1/2020). To investigate the advantage conferred by S:H655Y, we analyzed the kinetics, spike processing by cellular proteases and syncytium formation ability of a panel of SARS-CoV-2 variants harboring 655Y, including human isolates derived from patients seeking care at the Mount Sinai Health System in New York (NY) City which was one of the major early epicenters of COVID-19 pandemic. Our results demonstrated that the 655Y polymorphism enhances spike cleavage and viral growth. Furthermore, the S:655Y substitution was transmitted more efficiently than its ancestor S:655H in the hamster infection model. Finally, and in the context of the current epidemiological situation, we analyzed a set of emerging SARS-CoV-2 variants to investigate how different sets of mutations may impact spike processing. We demonstrated that novel circulating VOCs that became more prevalent have independently acquired mutations associated with a gain in spike cleavage and syncytia formation. Taken together, our study shows a link between an increased spike processing and increased virus transmission due to spike mutations present in SARS-CoV-2 variants that become epidemiologically more prevalent in humans.

## Results

### SARS-CoV-2 VARIANTS HARBORING 655Y SHOW AN ADVANTAGE IN REPLICATION AND ENHANCED SPIKE PROTEIN CLEAVAGE *IN VITRO*, AND IN TRANSMISSION *IN VIVO*

Minks have been suggested to play a role in the initial local spread and evolution of SARS-CoV-2 variants in different countries in Europe (*3, 4*). While minks are susceptible to SARS-CoV-2, they are also capable for zoonotic transmission of SARS-CoV-2 to because of the similarity of the ACE2 receptor between minks and humans. We used the mink model to investigate the replication and pathogenicity of the WA1 (USA-WA1/2020) isolate of SARS-CoV-2, as a representative of the first original human viruses that initiated the SARS-CoV-2 pandemic. This variant corresponds to one of the first USA isolates and does not contain any changes on the S protein when compared to the initial isolates from Wuhan, such as the Wuhan-1 virus. For this purpose, six minks were intranasally infected with 10^6^ pfu of WA1 isolate resulting in productive viral replication in the upper respiratory tract with infectious virus recovered from nasal washes at days 1, 3 and 5 post-infection (Supplementary Figure 1A-B). At day 4 post-inoculation, infectious virus was detected by plaque assays from left cranial lung and nasal turbinates but not from any of the other tissues analyzed (Supplementary Figure 1C). We then selected small and large viral plaques in the Vero-E6 cell-based plaque assays from infected mink lung specimens and performed next generation sequencing of the genome from the plaque-isolated viruses. As compared to the Wuhan-1 and WA1 reference sequences, all mink-derived viral isolates encoded the H655Y amino acid substitution within the spike (S) (Figure 1A). Additionally, the three viral isolates with the small plaque phenotype encoded the T259K amino acid substitution while the three viral isolates with the large plaque phenotype encoded the R682W amino acid substitution. It is known that S:682W/Q substitution in the furin cleavage site region may emerge after subsequent passages in VeroE6 cells (*29, 30*). Therefore, this mutation may have been selected during the course of the VeroE6 infections and not during the infection in minks. On the other hand, S:655Y appeared dominant in all the mink isolates, indicating that this mutation may confer an advantage in the mink host.

**Figure 1.**
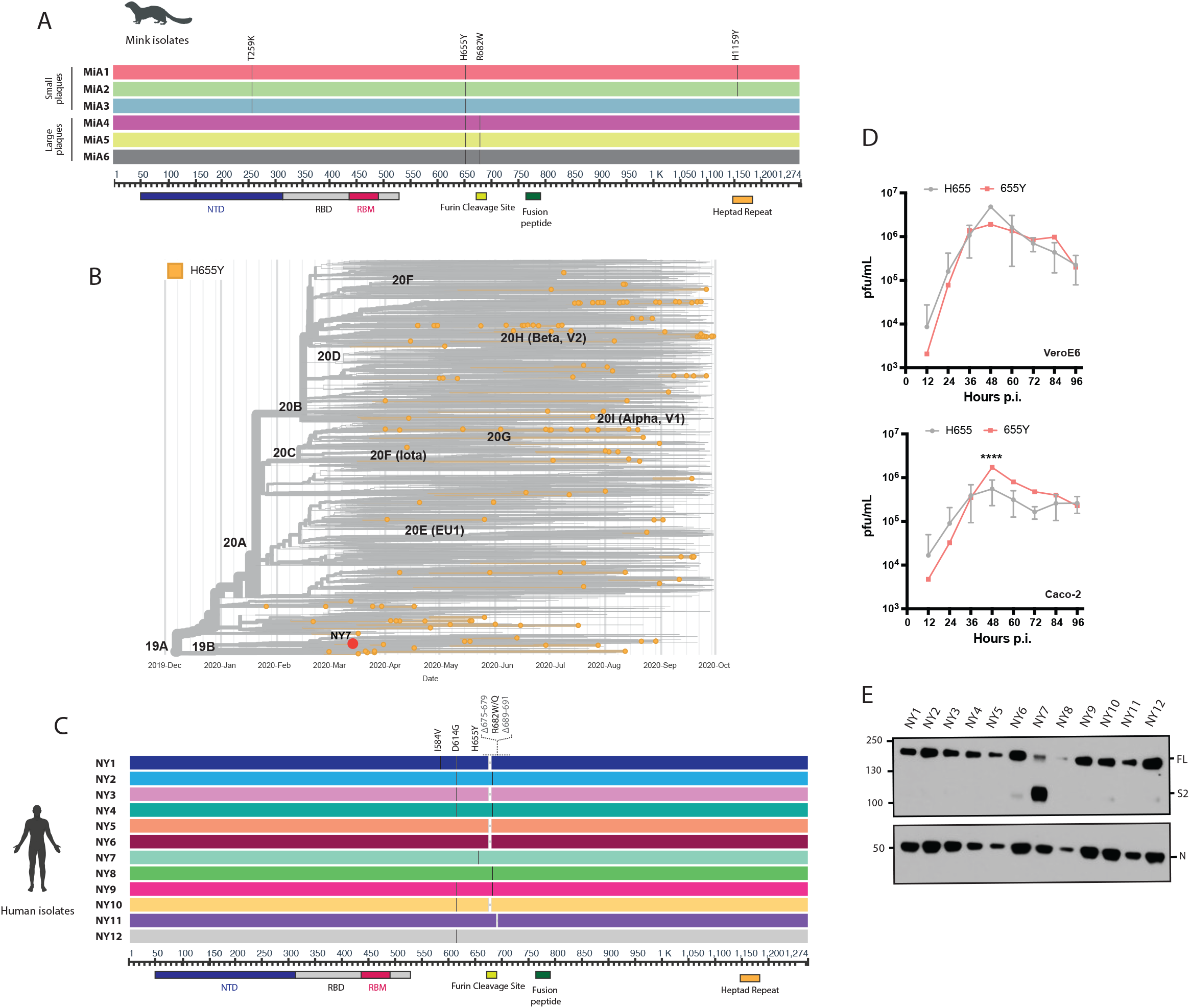
Mink and human SARS-CoV-2 variants bearing the 655Y polymorphism. A) Multiple sequence alignment of the spike (S) protein from SARS-CoV2 viruses isolated after infection in minks with WA1 isolate. Diagram shows the corresponding S amino acid substitutions mapped to the S gene. B) Time-calibrated phylogenetic analysis of the global distribution of H655Y substitution during the early SARS-CoV-2 outbreak. The phylogenetic tree was generated with Nextstrain with 7059 genomes sampled for representation of the H655Y substitution over time of worldwide data deposited in the GISAID database from December 2019 to September 2020. C) Multiple sequence alignment of the S protein from SARS-CoV2 viruses isolated from nasal swabs collected during the first pandemic wave in NY. Diagram shows the corresponding S amino acid substitutions mapped to the S gene. D) Viral growth of the NY7 containing the 655Y (red) versus its ancestors 655H (grey) in VeroE6 and Caco-2 cells. Cells were infected at an MOI of 0.01 and supernatants were titrated at the indicated hours post-infection (p.i.) and expressed as plaque forming units per milliliter (PFU). Means and SD are shown for the NY isolates containing 655H. ANOVA test was performed to compare mean differences within each group at different time points. Statistical significance was considered when p ≤ 0.05 (****, p < 0.0001). E) Western blotting of spike protein cleavage from supernatants of VeroE6 infected cells. Infections were performed at an MOI of 0.01 and supernatants were collected at 48 hours p.i. Full length (FL) spike protein (180 kDa), S2 cleaved spike (95 kDa) and Nucleocapsid (N, 50 kDa) were detected using specific antibodies. Levels of N protein were used as loading control.

To understand the magnitude and the spread of the 655Y polymorphism over time, we investigated the frequency of S:655Y over time in sequences sampled worldwide since the initial outbreak to the end of the first wave of SARS-CoV-2 (Figure 1B). For this, 7,059 sequences sampled from GISAID up to September 2020 were used. Human variants harboring the 655Y mutation were spread throughout the phylogenetic tree and distributed in all clades with no differences according to temporal distribution, suggesting that the 655Y mutation arose independently multiple times. Remarkably, the S:H655Y polymorphism was also found among the initial variants introduced in New York (NY) City in March 2020. To determine the replication phenotype, we decided to investigate this NY 655Y variant (NY7) together with some of its contemporaneous SARS-CoV-2 isolates circulating in New York during the early pandemic outbreak (*31*). To this end, we isolated 12 viruses based on their genotypes *(31)*, including NY7 which carries the S:655Y mutation for culture directly from nasopharyngeal specimens obtained from COVID-19 infected patients. Of note, the dominant 614G spike polymorphism was present in seven (58%) of the selected human SARS-CoV-2 (hCoV) NY isolates consistent with its early emergence and rapid spread worldwide (*17, 18*). Confirmation sequencing of the isolates showed that 682W/Q substitutions appeared in four (33%) viruses after initial isolation and culturing in VeroE6 cells. This is consistent with *in vitro* adaptative mutations previously described (*30*). Moreover, a five amino acid sequence (Δ675-679) flanking the furin cleavage site was deleted in five (42%) of the isolates as compared to the sequence from the original specimen. This deletion has been previously reported to be a common *in vitro* mutation selected in Vero cells (*29*). Amino acids substitutions of the S protein of these initial human isolates compared to the Wuhan-1 reference are shown Figure 1C. We next studied the replication kinetics of the NY SARS-CoV-2 isolates by comparing their multicycle growth curves at an MOI of 0.01 in VeroE6 and human Caco-2 cells. As expected, NY2 and NY4 containing the 682Q/W showed an advantage in VeroE6, while no differences could be found in Caco-2 cells (Supplementary Figure 2A-B). Remarkably, NY7 (S:655Y) showed higher growth at 48 hours post-infection (p.i) in Caco-2 cells (Figure 1D, Supplementary Figure 2B) when compared to the rest of these early SARS-CoV-2 isolates. These results support our conclusion that the 655Y polymorphism conferred a viral advantage. To investigate the spike cleavage efficiency of the 655Y versus other human isolates, we performed infections in VeroE6, and supernatants were analyzed by Western blot for the S2 domain of the S protein. Importantly, two bands were clearly visible for the NY7 (S:655Y) (Figure 1E), corresponding to both the cleaved (95 kDa) and uncleaved (180 kDa) form of the S protein. In contrast, only the uncleaved S form was detected in the other early human isolates indicating that the 655Y polymorphism may facilitate S protein processing.

**Figure 2.**
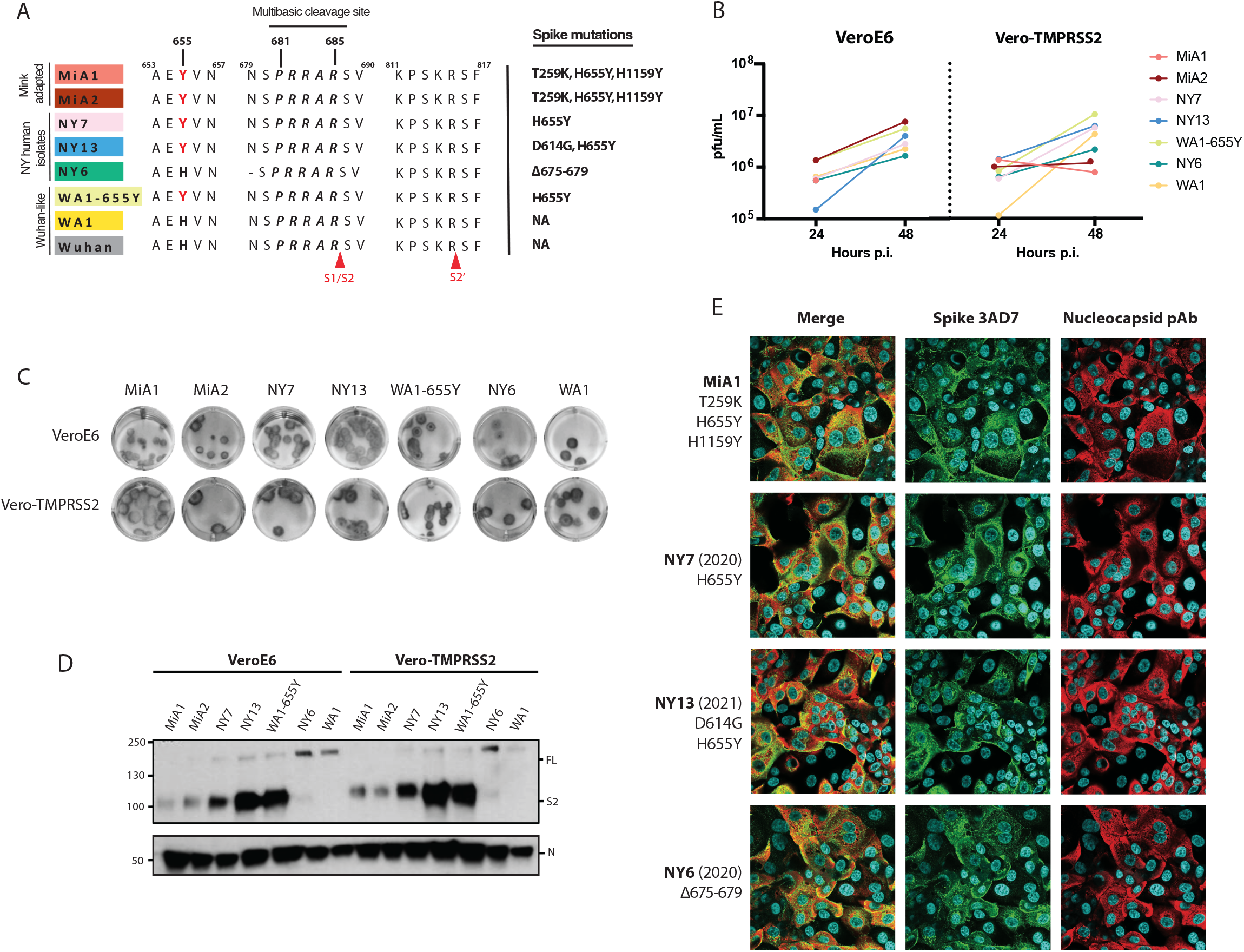
The H655Y amino acid substitution enhances spike cleavage and viral growth. A) Multiple alignment of the amino acid sequences around the 655 position and S1/S2 and S2’ cleavage sites from a panel of mink-adapted, cat, early human SARS-CoV-2 New York (NY) variants and WA1 virus. Wuhan1 is included as a reference. Red arrows indicate the location of S1/S2 and S2’ cleavage sites. Spike polymorphisms present across the S gene for each SARS-CoV-2 isolate are also shown. B) Replication kinetics of early SARS-CoV-2 viruses in Vero and Vero-TMPRSS2 cells. Infection was performed at an MOI of 0.01. Viral titers were determined by plaque assay at the indicated hours post-infection and expressed as PFU per milliliter. Color codes relate to the isolates shown in A. C) Plaque phenotype according to TMPRSS2 expression in Vero cells. The same viral supernatant was used to infect Vero and Vero-TMPRSS2 cells. Plaques were developed by immunostaining. D) Western blotting of S protein from supernatants of Vero and Vero-TMPRSS2 infected cells. Infections were performed at an MOI of 0.01 and viral supernatants were collected at 48 hours post-infection (p.i.). Full length (FL) spike protein (180 kDa), S2 cleaved spike (95 kDa) and Nucleocapsid (N, 50 kDa) were detected using specific antibodies. Levels of N protein were used as loading control. E) Immunofluorescence of SARS-CoV-2 S and N protein localization in Vero-TMPRSS2 infected cells at an MOI of 0.01 and 24 hours p.i. Spike protein was detected using a specific monoclonal antibody 3AD7 (green), N protein was detected using a polyclonal antiserum (red) and 4’,6-diamidino-2-phenylindole (DAPI) was used to stain the nucleus.

To confirm whether the 655Y mutation was solely responsible for the increased S cleavage, we analyzed the replication and cleavage efficiency of a panel of SARS-CoV-2 viruses, all bearing the 655Y substitution but containing additional substitutions across the genome. We included two of the isolated mink variants (MiA1 and MiA2); NY7 and another human isolate derived from a COVID patient infected in February 2021 (NY13, S:614G, 655Y); and a previously published WA1-655Y variant isolated after wild type WA1 infection in cats (*32*). Additionally, the WA1 reference and NY6 were used as controls since they lack the 655Y substitution. It should be noted that NY6 has a five amino acid deletion before the furin cleavage site (Figure 1C and 2A). We assessed differences in replication and S processing of this panel of viruses by comparing growth in both VeroE6 and Vero-TMPRSS2 cells. As shown in Figure 2B, WA1-655Y infection yielded higher titers in both VeroE6 and Vero-TMPRSS2 cells as compared to infection by WA1. These isolates only differ in the position 655Y while the rest of the genome is isogenic, supporting that 655Y spike polymorphism enhances viral replication and growth. Next, viral supernatants were used to analyze the plaque phenotype in VeroE6 and Vero-TMPRSS2 and to compare S protein expression levels after infection. In general, all isolates showed higher plaque size in the presence of TMPRSS2, consistent with enhancement of cell entry (Figure 2C). However, differences were found by Western blot and only the isolates bearing 655Y showed enhanced spike cleavage in both VeroE6 and Vero-TMPRSS2 (Figure 2D). Finally, and to investigate the ability to induce syncytia, we infected Vero-TMPRSS2 at an MOI of 0.01 with the mink (MiA1) and human isolates (NY7, NY13 and NY6) and used specific antibodies to detect the S and N protein, as well as nuclei staining with DAPI after 24 p.i. by immunofluorescence microscopy. As shown in Figure 2E, the isolates harboring the 655Y polymorphism showed a slight increase in syncytia formation as compared to NY6 (S:Δ675-679).

Syrian golden hamsters are a recognized rodent model to investigate infection and transmission of SARS-CoV-2 (*33, 34*). To test whether the S:655Y polymorphism enhances viral transmission, five pairs of Syrian golden hamsters were placed in individual cages to perform viral competition experiments. For this, one hamster of each pair was infected intranasally with 10^5^ pfu of a mix of WA1 and WA1-655Y viruses at a one-to-one ratio (Figure 3A). Direct infected (DI) and direct contact (DC) hamsters were euthanized after day 5 and 7 post-infection, respectively, and lungs and nasal turbinates were harvested for subsequent viral titer quantification. In addition, nasal washes were collected on day 2 and 4 in both DI and DC, and on day 6 p.i. of DC animals. One of the DI hamsters died after nasal wash collection at day 2, leaving 4 animals subjected to follow up in the DI group. Hamsters were also monitored daily for body weight loss. After 2 days p.i. DC hamsters exhibited a decrease in weight indicating early viral transmission from infected animals (Figure 3B). This observation was further supported by detection of infectious virus in nasal washes at 2 days p.i in both DI and DC hamsters. At day 4 p.i., viral titers were detected in 3 out of 4 animals and at day 6 p.i., viral replication was not detected in two of the DC nasal wash samples (Figure 3C). In general, we observed a decrease over time in the infectious virus present in nasal washes from DI and DC animals. We then determined the relative abundance of S:655Y on the viral RNA present in the nasal washes by next-generation sequencing (Figure 3D). The consensus RNA sequence from all DI hamsters contained the S:655Y polymorphism suggesting that WA1-655Y virus was able to overcome the wild type WA1 isolate during infection. Similarly, S:655Y was present in all the nasal washes collected from four DC hamsters indicating an advantage conferred by this mutation in viral transmission. However, one DC animal (hamster 4-C) showed a decrease of 655Y abundance over time. Interestingly, this hamster lost less weight when compared to the rest of the animals. Next, we analyzed the viral growth in lungs and nasal turbinates from DI (collected at day 5 p.i.) and DC (collected at day 7p.i.) hamsters (Figure 3E). No differences were found in viral titers in the tissues from both animal groups. However, we observed that three DC hamsters had lower lung titers compared to the rest of the animals. These same hamsters also exhibited low viral loads in the nasal turbinates. We then sequenced the viral RNA present in these tissues (Figure 3F-G). The RNA from one lung and nasal turbinate of one DI and DC hamster could not be amplified by specific PCR for downstream sequencing. Figure 3F-G shows that all lungs and nasal turbinate tissues from DI animals analyzed had the S:655Y mutation. In addition, S:655Y was present in 75% of the nasal turbinates and lungs from DC animals. Taken together, our results demonstrate that 655Y polymorphism increases spike cleavage and viral growth. Moreover, viral competition and transmission experiments showed that S:655Y became predominant in both DI and DC animals indicating that this mutation plays a role in viral transmissibility.

**Figure 3.**
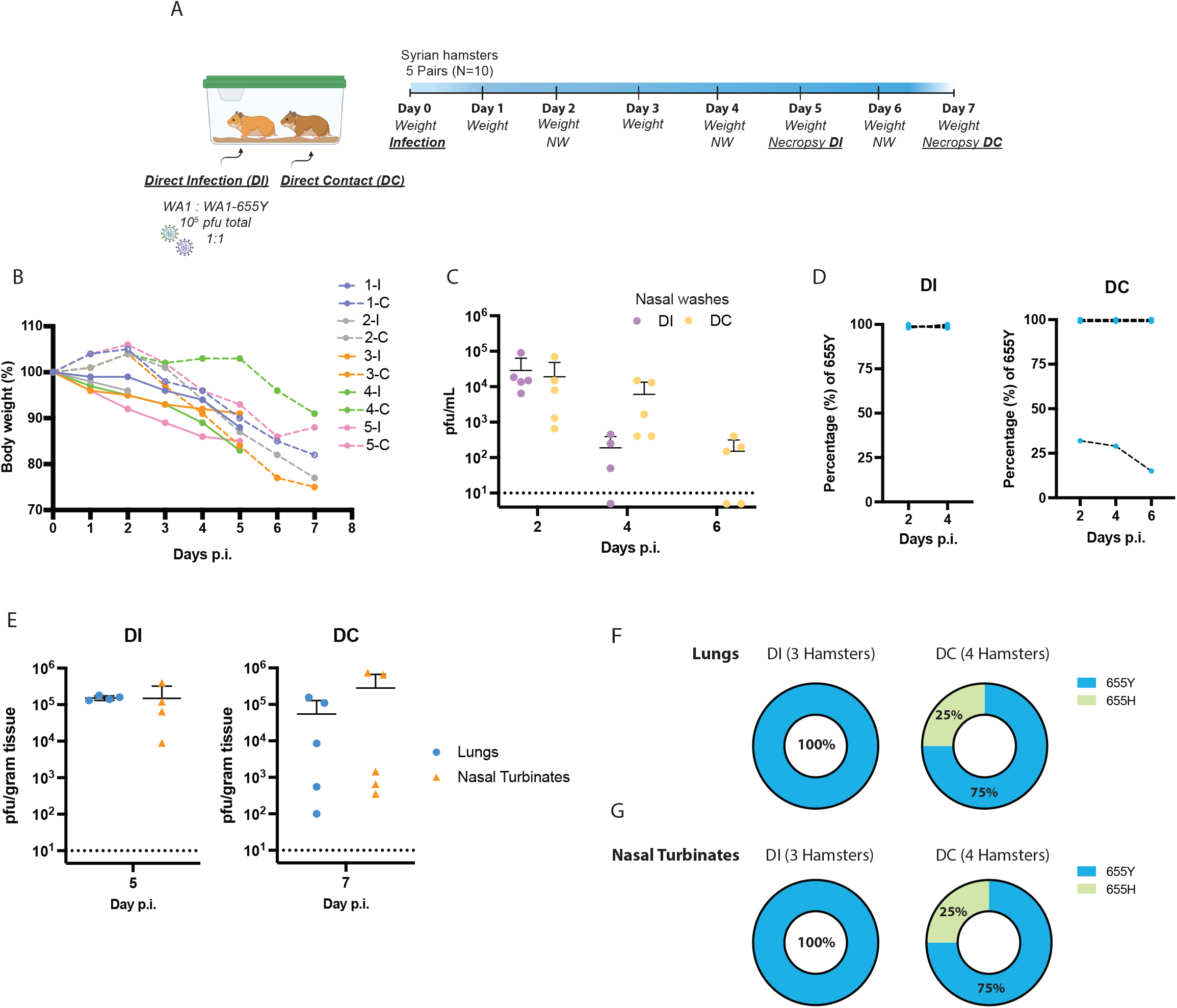
The 655Y polymorphism prevails over the 655H in the transmission *in vivo* model. A)Ten 3-weeks-old female Syrian hamsters were placed in pairs. Only one hamster per cage was infected intranasally with a total of 10 ^5^ pfu of SARS-CoV-2 WA1 and WA1-655Y isolates in a one-to-one ratio. Nasal washes were collected at day 2, 4 and 6 post-infection (p.i.). Lungs and nasal turbinates were harvested from direct-infected (DI) and direct-contact (DC) hamsters at day 5 and 7 p.i., respectively. B) Body weight change of individual hamsters over time. C) Viral titers of nasal washes expressed as PFU per milliliter. Error bars indicate SDs. D) Relative abundance of 655Y mutation in the RNA from nasal washes in the DI and DC hamsters. The y axis shows the percentage of 655Y polymorphism in the total good quality sequencing reads from each biological RNA sample and the x axis indicates the day p.i. samples were collected. E) Viral titers of lungs and nasal turbinates expressed as PFU per gram of tissue. Error bars indicate SDs. Titers of DI and DC hamsters are shown at day 5 and 7 p.i., respectively. F-G) Proportion of hamsters with 655Y (blue) and H (green) in the nasal turbinates and lungs from DI and DC as confirmed by next generation sequencing.

### SARS-CoV-2 VARIANTS EVOLVE TO ACQUIRE AN INCREASED SPIKE CLEAVAGE AND FUSOGENIC ABILITY

Current circulating SARS-CoV-2 VOCs bear novel spike polymorphisms that correlate with an enhanced human transmission (*19*) and reduced antibody neutralization (*22, 35*). Interestingly, the Gamma variant (lineage P.1) which emerged in Brazil in November 2020, harbors the amino acid spike substitution H655Y. In the context of the evolving epidemiological situation, we decided to investigate whether the co-emergence of different selective mutations in some representative VOC had a similar phenotype to SARS-CoV-2 viruses harboring the S:655Y. We first estimated the amino acid substitution frequencies around the cleavage site region (655 to 701) from globally available data (2,072,987 sequences deposited in GISAID Database up to 28 June 2021). As expected, the P681H/R, H655Y and A701V substitutions showed high prevalence since they are harbored by the main VOCs lineages. Additionally, the less prevalent Q675H and Q677H were also found to be a highly variable position and present in widely circulating variants (*36*). We then spatially and structurally mapped these amino acid changes within and surrounding the furin cleavage sequence of the S protein (Figure 4A). The 655 position was located in close proximity to the furin cleavage site. Next, we performed a phylogenetic analysis of sequences sampled worldwide from February 2020 to June 2021 to illustrate the temporal distribution and phylogenetic relationship of the high prevalent S mutations (Figure 4B). For this, a sample set of 13,847 sequences deposited in GISAID up to June 2021 were analyzed. While the H665Y frequency was higher in the Gamma lineage (P.1), it could be found also in 19B clade (Figure 4C), in line with our identified NY7 isolate. The P681H substitution located in the furin cleavage site of the spike protein was identified in the Alpha variant that emerged in September 2020. Interesting, this mutation was also found in the Theta variant, first detected in February 2021 (Figure 4D). In contrast, Kappa and Delta variants harbor polymorphism P681R (Figure 4D). Finally, the frequency of A701V mutation was higher in Beta and Iota variants which emerged in October and November 2020, respectively (Figure 4E).

**Figure 4.**
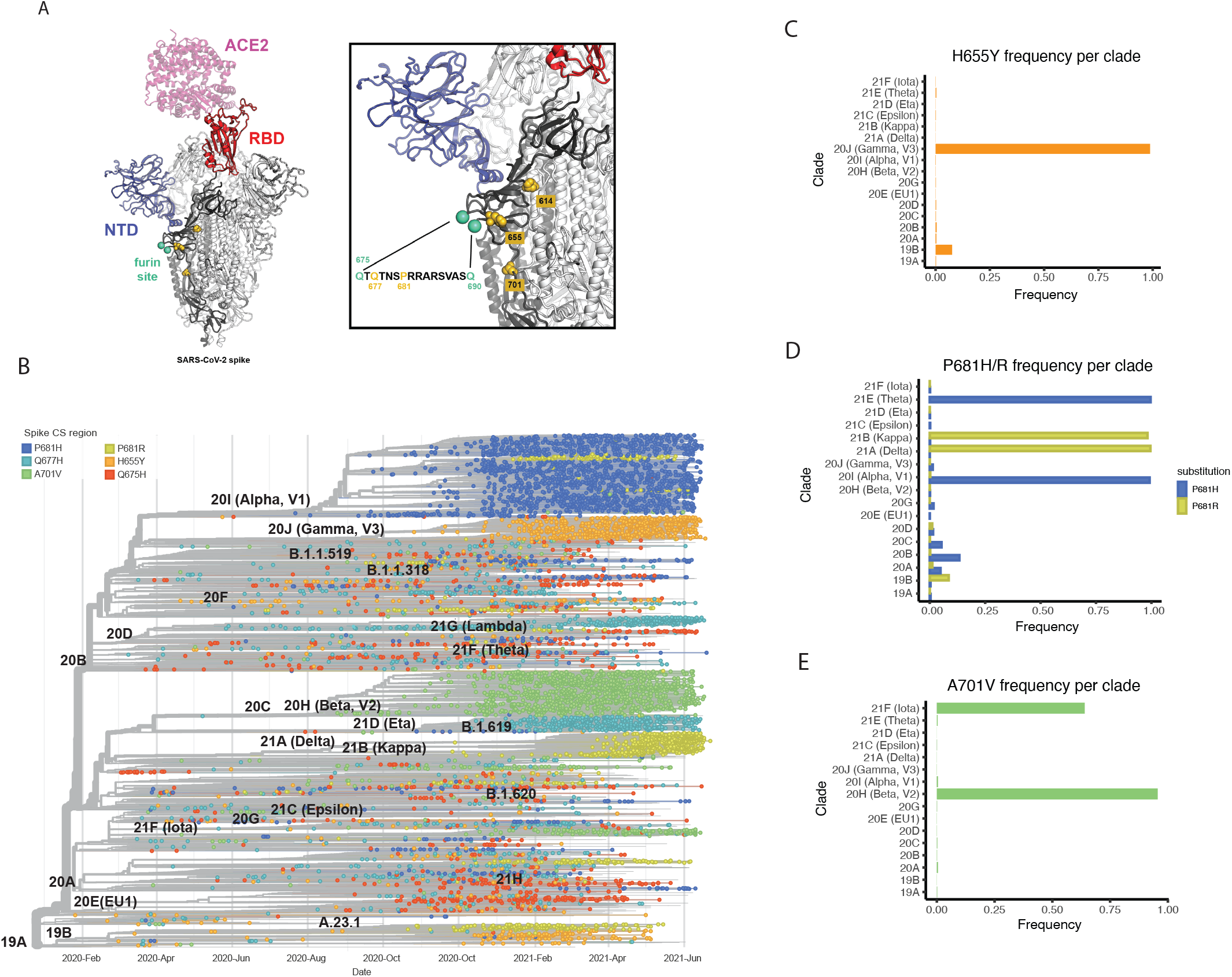
Global epidemiology of SARS-CoV-2 variants of concern (VOCs). The amino acid substitution frequencies around the cleavage site region (655 to 701) from globally available data (2,072,987 sequences deposited in GISAID Database as of 28 June 2021) was estimated. A) Shows the high prevalent mutations identified mapped onto the structure of the S glycoprotein. The model was generated by superposition of PDB 6M0J and 7C2L (*40, 41*). One RBD in the up conformation (red) is bound with ACE2 receptor (pink). The NTD is colored blue, the amino-acid substitutions are shown as gold spheres and the furin cleavage loop (disordered and therefore missing in most atomic models) is flanked with cyan spheres. One spike protomer is shown in bold colors while the other two are colored white. A zoomed-in image of the region of interest and the sequence of the furin site loop is also shown. Amino-acid residues of interest are highlighted in gold. B) Time-calibrated phylogenetic tree of SARS-CoV-2 circulating variants illustrating the temporal distribution and phylogenetic relationships of the most prevalent S mutations along the S1/S2 region (highlighted in color). The phylogenetic tree was generated using NextStrain and analysis was performed using a sample of 13,847 genomes focused on the most prevalent substitutions between S:655 and S:701 between February 2020 and June 2021 from GISAID database. C-E) Frequency per clade of H655Y, P681H/R and A701V spike polymorphisms.

Next, we analyzed the *in vitro* phenotype of some of the most prevalent SARS-CoV-2 VOCs. Multiple protein sequence alignment of the VOCs used are shown in Figure 5A. In contrast to the NY isolates previously analyzed, VOCs showed a considerable number of unique changes across the whole spike protein. However, all of them showed similar replication kinetics and plaque phenotypes in Vero-TMPRSS2 cell monolayers. Conversely, clear differences were observed when replication efficiency was determined in VeroE6 cells at 24 hours post-infection (p.i.) (Figure 5B). Additionally, viral supernatants were titrated on both VeroE6 and Vero-TMRPSS2 cells (Figure 5C). Substantial differences in plaque phenotypes were observed, especially for the Kappa (B.1.617.1) and Delta (B.1.617.2) variants, when TMRPSS2 was present. Thus, these late VOCs that emerged late in the COVID-19 pandemic might strongly depend on the presence of TMRPSS2 to establish optimal infection *in vitro*. Last, we investigated the extent of the spike cleavage of these VOCs. For this, infections were performed in VeroE6 and Vero-TMPRSS2 at an MOI of 0.01 and viral supernatants were analyzed after 48 hours post-infection (p.i.) by Western blot (Figure 5D). N protein was used as a control for viral replication and loading. Similarly, WA1 was included as a reference since no selective mutations are found in the S protein. Figure 5D shows similar cleaved and uncleaved S protein levels for all the SARS-CoV-2 VOCs in the presence of TMRPSS2 expressed in Vero-TMPRSS2 cells. In contrast, only Beta (B.1.351) and Gamma (P.1), exhibited an increased spike cleavage when the infections were performed in wild type VeroE6 cells. Interestingly, the spike and nucleocapsid expression of Kappa (B.1.617.1) and Delta (B.1.617.2) variants was not detectable by Western blot analysis of VeroE6 supernatants. The canonical cleavage at the S1/S2 site occurs at the last arginine (R) of the multibasic PRRAR motif and is performed by furin proteases at this specific residue. Thus, we next quantified the abundance of furin-cleaved peptide of VOCs in Vero-TMPRSS2 cell supernatants by targeted mass spectrometry. Vero-TMPRSS2 cells were infected at an MOI of 0.1 with the indicated VOCs and NY7 (S:H655Y) and WA1-655Y isolates. WA1 and NY6 were used as controls. Cell extracts were collected after 24 hours post-infection and samples were prepared. The abundance of the C-terminal peptide resulting from endogenous furin cleavage at the terminal arginine (PRRAR \ SVASQSIIAYTMSLGAE) was quantified as a proxy of cleavage efficiency since this peptide is common for all SARS-CoV-2 VOCs, except for the Beta that contained a V instead of A at the end of the peptide (SVASQSIIAYTMSLGVE). Fold change peptide-level abundance for each variant compared to WA1 control was calculated and plotted in Figure 5E. Isolates WA1-655Y, NY7 and Gamma, all of them harboring the 655Y mutation, and Beta harboring 701V, showed the higher abundance of furin-cleaved peptide. Conversely, lower levels of C-terminal cleaved peptide were found for the VOCs harboring the 681H/R amino acid change suggesting that an introduction of an amino acid change at this position might modify the canonical cleavage residue at the last R of the furin cleavage site. Nonetheless, when we assessed the fusogenic capacity of the S protein of VOCs by immunofluorescence microscopy of infected Vero-TMPRSS2 cells, we found strong syncytium formation induced by all the variants (Figure 5F). Interestingly, extensive fusogenic capacity was also exhibited by Delta and Kappa variants (Figure 5F-G) consistent with abundant cleaved S form found by Western blot (Figure 5D). Because cleavage at the multibasic furin motif is believed to be required for optimal syncytium formation (*10*), we finally compared the ability to induce cell fusion by the Kappa variant and a mutated form lacking amino acids at the furin cleavage site (S: Δ678-682). This Δ678-682 Kappa was obtained after consecutive passage and culturing in VeroE6 cells. As shown in Figure 5G, a loss of fusogenic activity was observed when compared to the intact Kappa VOC. Altogether, our results are consistent with the notion that current highly transmissible circulating VOCs have evolved independently to acquire mutations associated with increased spike protein processing and transmission.

**Figure 5.**
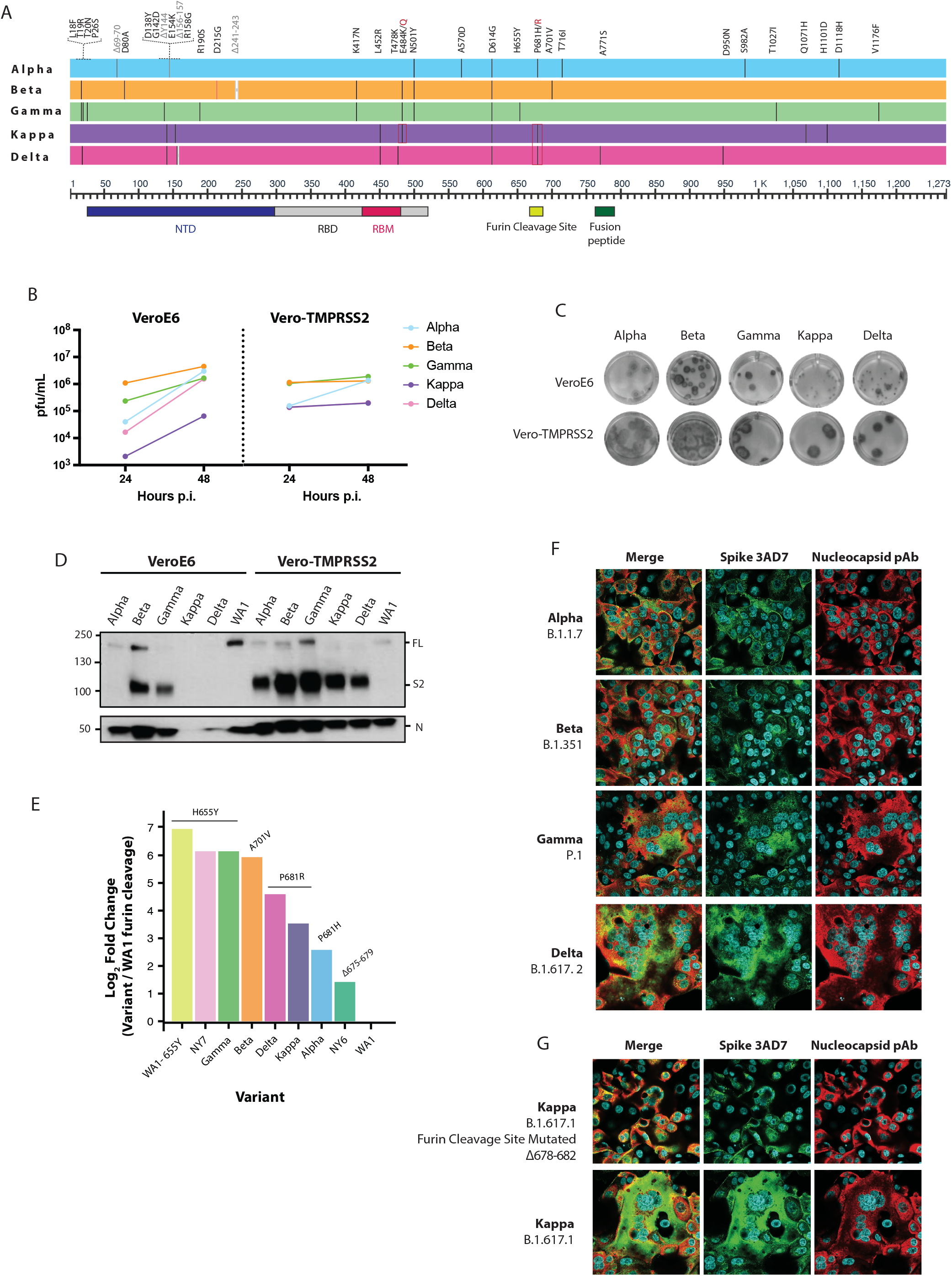
SARS-CoV-2 VOCs evolve to a convergent phenotype associated to an increase on S cleavage and fusogenicity. A) Multiple alignments of the S protein of the indicated SARS-CoV-2 VOCs. Diagram shows the corresponding S amino acid substitutions mapped to the S gene. Viral growth of SARS-CoV-2 variants in Vero and Vero-TMPRSS2 cells. Infections were performed at an MOI of 0.01. Viral titers were determined by plaque assay at the indicated hours post-infection and expressed as PFU per milliliter. C) Plaque phenotype of VOCs according to TMPRSS2 expression. Same viral supernatant was used to infect Vero and Vero-TMPRSS2 cells. Plaques were developed by immunostaining. D) Western blotting of spike cleavage in supernatants from Vero and Vero-TMPRSS2 infected cells at an MOI of 0.01. Viral supernatants were collected at 48 hours post-infection. Full length (FL) S protein (180 kDa), S2 cleaved spike (95 kDa) and Nucleocapsid (N, 50 kDa) were detected using specific antibodies. Levels of N protein were used as loading control. E) Quantification of the clevage efficiency by mass spectrometry. Vero-TMPRSS2 cells were infected at an MOI of 0.1 with the indicated VOCs and NY7 (S:H655Y) and WA1-655Y isolates. WA1 and NY6 were used as controls. Cells extracts were collected after 24 hours post-infection. Cleavage efficiency was determined by measuring the abundance of the resulting peptide (<u>SVASQSIIAYTMSLGAE)</u> after cleavage at the terminal arginine of the furin cleavage site. The y axis shows the log_2_ of fold change between cleaved peptide abundance for each variant normalized by WA1 control. F-G) Immunofluorescence of SARS-CoV-2 S and N protein in Vero-TMPRSS2 infected cells at an MOI of 0.01 and 24 hours p.i. for the indicated VOCs. Spike protein was detected using a specific monoclonal antibody 3AD7 (green), N protein was detected using a polyclonal antiserum (red) and 4’,6-diamidino-2-phenylindole (DAPI) was used to stain the nucleus.

## Discussion

Emerging SARS-CoV-2 VOCs contain novel spike polymorphisms with unclear functional consequences on epidemiology, viral fitness, and antigenicity. In this study, we evaluated the impact of different spike mutations on viral infection, pathogenicity, and *in vivo* transmission. We found that in the mink animal model the 655Y spike substitution is selected after infection with the WA1 isolate. Phylogenetic analysis of genome sequences collected worldwide showed an early sporadic appearance of S:655Y during the first pandemic wave in New York in March 2020, and the presence of this mutation in several posterior lineages, including SARS-CoV-2 Gamma variant, pointing to a potential role in adaptation and evolution. To better understand the impact of this polymorphism, we isolated and *in vitro* characterized a panel of SARS-CoV-2 viruses bearing the 655Y spike mutation. Our results demonstrated that S:655Y enhances the viral growth and the spike protein processing required for optimal cell entry and viral-host membrane fusion. In addition, we performed viral competition and transmission experiments in the hamster animal model and showed that S:655Y became predominant in both direct infected and direct contact animals. Finally, we showed that VOCs converge to gain spike cleavage efficiency and fusogenic potential.

Here, we demonstrate that viruses containing the H655Y polymorphism confer a growth advantage in both VeroE6 and human-like Vero-TMPRSS2 cells. Interestingly, the early human isolate NY7 harboring the 655Y mutation also showed higher replication in human Caco-2 cells. However, it is known that other mutations outside of the S gene could be impacting viral replication and infection (*37, 38*). Therefore, we confirmed the S:655Y mutation alone was responsible for the enhanced growth and spike cleavage phenotype when comparing WA1 wild type and WA1-655Y isolates. These variants have the same viral protein amino acid sequence except for the amino acid present at position 655 of the spike. Since most of the isolates used in this study contain a constellation of mutations across the genome that could increase viral fitness, comparison of both viruses in parallel allowed to detect differences in growth and spike cleavage that can be attributed only to 655Y polymorphism. S:655Y is present in the S1 spike domain outside of the RBD and has been associated with a decrease of the neutralizing activity when targeted by some monoclonal antibodies(*26*). However, H655Y has been also naturally selected in cats and mice suggesting a beneficial impact of this substitution in widen viral host range and susceptibility (*27, 28*). Our data further supports this argument because we also found that S:655Y is selected after replication in minks, a natural host for SARS-CoV-2. Besides, when we assessed the viral transmission efficiency of 655Y versus the ancestor 655H in competition experiments in the hamster model, we also found that 655Y becomes more prevalent, as the bulk of infectious viruses recovered from the infected animals harbored this mutation, except for one hamster. This indicates that S:655Y can overcome S:655H *in vivo*.

Intense worldwide surveillance has established that SARS-CoV-2 variants are constantly emerging. In particular, the spike protein has shown high plasticity (*6*). Most of the spike mutations associated with a decrease in neutralization by antibodies against earlier viruses are located in the RBD or N-terminal domain (NTD), which are critical for binding and interacting with the ACE2 cellular receptor. While mutations at these domains may impact SARS-CoV-2 vaccine efficacy, it is also vital to characterize other mutations that might explain the gain in transmissibility observed for the VOCs. Since the Gamma variant that emerged in November 2020 also harbors the 655Y polymorphism (Figure 5A), we decided to investigate its phenotype *in vitro*. Similar to the earlier S:655Y isolates, this variant also exhibited an increase in spike processing efficiency. More importantly, this phenotype was also confirmed in all emerging VOCs analyzed when infections were performed in the Vero-TMPRSS2 cells indicating that additional mutations within S confer this advantage. Most likely, the spike mutations P681H in Alpha variant-first identified in United Kingdom-and P681R harbored by Kappa and Delta variants-first emerged in India-allowed this enhanced S cleavage. Interestingly, for these variants, optimal cleavage appeared to be dependent on TMPRSS2 protease activity (Figure 5D).

To confirm the cleavage at the putative furin cleavage site, we determined the relative abundance of the furin cleaved peptide produced after the 685-terminal arginine. We observed higher amount of cleavage at this position as compared to the previous circulating viruses, although lower amounts were detected in Alpha, Kappa and Delta variants as compared to the viruses harboring the 655Y mutation. This suggests that a change in residue 681 may introduce an additional cleavage site, perhaps recognized by TMPRSS2 protease that enhances spike cleavage of these variants and produces an additional cleavage peptide different in size and amino acid sequences. Further research is needed to confirm the existence of a recognition site for additional proteases different than furin in the amino acid motif SH/RRRAR when the P681S/H mutation is present. In any case, all the VOCs analyzed proved to be strong syncytia inducers which could potentially indicate a role in pathogenesis and lung damage mediated by TMPRSS2 activity after infection in humans (*39*). On the other hand, the Beta variant, which was first identified in South Africa in October 2020, does not contain a change in the furin cleavage site or in the spike position 655, but instead a change in the residue found at position 701. Although this residue is found around 20 amino acids away from the furin cleavage motif, we found similar results when the extent of the spike processing was investigated (Figure 4A-E; 5D-G). It is important to note that the VOCs investigated in here independently acquired S mutations around the furin cleavage site that became epidemiologically more prevalent in humans. When we investigated the spatial distribution by superimposition of the crystal structure of the S protein, we found that these highly prevalent polymorphisms were all located in close proximity to the furin site loop (Figure 4A). Any substitution in this protein domain is likely to have an effect on the structural integrity and dynamics, potentially impacting the accessibility of the polybasic site to the relevant protease and likely facilitating the recognition by furin.

In summary, our study demonstrates that the 655Y spike polymorphism, present in the Gamma VOC, is a key determinant of SARS-CoV-2 infection and transmission. The selection and increasing frequency of S:655Y in the human population and following SARS-CoV-2 infection of different animal models such as cats, mice and minks suggests this mutation is associated with an improvement of viral fitness and adaptation to diverse hosts through an increased cleavage of the spike protein. Additionally, we provide evidence of adaptative mutations that SARS-CoV-2 VOCs have been acquired and are associated with an increased spike protein processing. This has significant implications in the understanding of the viral determinants that can impact viral transmissibility, viral evolution, and possibly SARS-CoV-2 antigenicity and pathogenicity.

## Acknowledgements

We thank Richard Cadagan for excellent technical assistance. We also thank Florian Krammer for providing the monoclonal mouse anti-spike KL-S-3A7 antibody used in this study and Andy Pekosz and Mehul S. Suthar for providing Beta and Kappa VOCs, respectively. We also want to thank Lisa Miorin and Michael Schotsaert for training and support.

## Funding

This research was partly funded by CRIPT (Center for Research on Influenza Pathogenesis and Transmission), a NIAID funded Center of Excellence for Influenza Research and Response (CEIRR, contract #75N93021C00014) (AGS)

NCI SeroNet grant U54CA260560 (AGS)

NIAID grants U19AI135972 and U19AI142733 (AGS)

DARPA grant HR0011-19-2-0020 (AGS)

JPB Foundation (AGS)

Open Philanthropy Project (research grant 2020-215611 (5384) (AGS)

Anonymous donors to AGS

AGR is funded by Marion Alban MSCIC Scholars Award and the 2020 Robin Chemers Neustein Postdoctoral fellowship

ML is funded by a fellowship of the Belgian American Education Foundation

NBAF Transition Funds from the State of Kansas (JAR)

NIAID Centers of Excellence for Influenza Research and Surveillance under contract number HHSN 272201400006C (JAR)

AMP Core of the Center for Emerging and Zoonotic Infectious Diseases of the National Institute of General Medical Sciences (NIGMS) of the National Institutes of Health under award number P20GM130448 (JAR)

Department of Homeland Security Center of Excellence for Emerging and Zoonotic Animal Diseases under grant number HSHQDC 16-A-B0006 (JAR).

## Author contributions

TA and AGS conceived, designed, and supervised the study. TA provided training to AE. AE performed most of the experiments including growth of viral stocks, viral infections and titration of growth curves, plaque phenotype analysis, sample preparation for western blot and mass spectrometry analysis, western blot experiments, RT-qPCR and infectivity analysis, immunofluorescence with the help of TA (viral isolation, plaque assay and plaque purification of viruses, immunofluorescence imaging acquisition), SA (titration by plaque assay analysis) and ML (immunofluorescence staining). AGR and AVG performed experiments for whole-genome and spike-specific sequencing. AGR performed genome assembly, variant calling, global phylogenetic and variant-prevalence analyses. IM provided Gamma VOC viral stock and performed the hamster competition experiment with the help of RP, ML and AE. RA and RP performed the mink infections, titration by plaque assay and mink-adapted S:655Y (MiA) viral purification and isolation experiments. AF performed the targeted pro teomics and analyzed the targeted proteomics data. RR provided Alpha and Beta VOCs viral stocks. TK provided mink selected (MiA1 and MiA2) viral stocks. DAM, VB, CM, and JAR isolated and provided the WA1-655Y cat selected-variant. VS and EMS provided human nasopharyngeal swabs from SARS-CoV-2 infected individuals. HA isolated SARS-CoV-2 viruses from human nasopharyngeal swabs. GB provided the spike structural analysis figure. MB, LZA, NK and HVB provided methods and expertise. AE and TA analyzed data, wrote the manuscript, and prepared the figures. All the authors reviewed and edited the manuscript.

## Data availability

All data is available in the manuscript or the supplementary materials. The targeted mass spectrometry data including raw files, Skyline document and resulting transition lists via ProteomeXchange with identifier PXD027641. Reagents used are almost exclusively commercially available and non-proprietary.

## Materials and Methods

### Cell lines

VeroE6 and Caco-2 cell lines were originally purchased from the American Type Culture Collection (ATCC). VeroE6-TMPRSS2 cell line was purchased from BPS Bioscience. A master cell bank was created for each cell line and early-passage cells were thawed in every experimental step. VeroE6 and Caco-2 cell lines were maintained in Dulbecco’s modified Eagle’s medium (DMEM) with glucose, L-glutamine, and sodium pyruvate (Corning) supplemented with 10% fetal bovine serum (FBS, Gibco), non-essential amino acids, penicillin (100 UI/mL), streptomycin (100 UI/mL) (Corning) and normocin (100 ug/mL) (InvivoGen) to prevent mycoplasma infection. VeroE6-TMPRSS2 cell line was cultured in the same growth media described above and further supplemented with sodium pyruvate (Corning) and puromycin (3 ug/mL) (InvivoGen). All cell lines were grown at 37°C in 5% CO_2_.

### Viruses

All experiments in this study were performed in the BSL-3 facility following Icahn School of Medicine biosafety guidelines. Human SARS-CoV-2: nasopharyngeal swab specimens were collected as part of the routine SARS-CoV-2 surveillance conducted by the Mount Sinai Pathogen Surveillance program (IRB approved, HS#13-00981). Specimens were selected for viral culture on Vero-E6 cells based on the complete viral genome sequence information (*31*). Human isolates NY1 to NY12 were obtained from the nasopharyngeal swabs of patients infected with SARS-CoV-2 in March 2020 while NY13 (PV28021) was cultured from the nasopharyngeal swab of a patient infected in February 2021. All nasal swabs were kindly provided by Dr. Viviana Simon. GISAID accession numbers of these isolates are shown in Supplementary Table 1. Viral transport media was incubated with VeroE6 cells until cytopathic effect was observed and supernatants from infected cells were used for plaque purification of clonal population as previously described (*42*). SARS-CoV-2 isolates USA-WA1/2020 (NR-52281), hCoV-19/England/204820464/2020 (NR-54000, Alpha) and hCoV-19/Japan/TY7-503/2021 (NR-54982, Gamma) were obtained from BEI resources. hCoV-19/USA/MD-HP01542/2021 JHU (Beta variant) was a gift from Dr. Andy Pekosz. Kappa and Delta variants were kindly provided by Dr. Mehul S. Suthar and Dr. Viviana Simon, respectively. Animal SARS-CoV-2: Mink SARS-CoV-2 variants (MiA-1 and MiA-2) were isolated during the mink experiment described below. After plaque assay analysis of the left cranial lung collected from a WA1 infected mink at 4 days p.i., two plaque phenotypes were observed. Only the small plaque phenotype viruses were grown and used in this study. USA-WA1/2020-H655Y (WA1-655Y) was kindly provided by Dr. Jurgen Richt. This variant was isolated after a WA1 cat infection. All the viral stocks were produced by infecting VeroE6 or VeroE6-TMPRSS2 cells at a MOI of 0.01. Infected cells were maintained in infection media (DMEM with glucose, L-glutamine, and sodium pyruvate supplemented with 2% FBS, non-essential amino acids, HEPES, penicillin (100 UI/mL) and streptomycin (100 UI/mL)) at 37°C in 5% CO_2_. Infected cells were monitored by microscopy and cell-infected supernatants were collected at day 2 post-infection when cytopathic effect was observed. Viral supernatants were clarified of cell debris by spin down followed by centrifugation at 2000 x g for 20 min in Amicon Ultra-15 centrifugal filters (Sigma, 100 kDa cutoff) to concentrate the viral stocks. Aliquots were stored at −80°C until titration by plaque assay. All SARS-CoV-2 variants were sequence-confirmed before performing the experiments.

### Infection of cell cultures

Approximately 3.2 × 10^5^ VeroE6 or VeroE6-TMPRSS2 or Caco-2 were seeded in a 12 well-plate and cultured at 37°C in 5% CO_2_ for 16 hours. Cells were infected with the corresponding SARS-CoV-2 isolate at an MOI of 0.01. Cells were incubated with the virus for 1 hour and then, cells were washed with PBS to ensure removal of non-attached virus. After infection, cells were maintained in infection media. Supernatants were collected at and the indicated time points and stored at −80°C for plaque assay analysis and virus quantification.

### Western blotting

VeroE6 or VeroE6-TMPRSS2 cells were infected with the indicated SARS-CoV-2 isolates, similar to the description above. Viral supernatants were collected at 24- and 48-hours post-infection. Supernatants were clarified by low-speed spin. Viral supernatants and cell extracts were mixed with RIPA buffer (Sigma Aldrich) containing EDTA-free protease inhibitor cocktail (Roche) and 10% SDS (Invitrogen) to a final concentration of 1%. Then, samples were boiled for 10 minutes at 100°C and centrifuged for 10 minutes at 4**°C** and maximum speed. Viral supernatants were subjected to SDS-PAGE protein electrophoresis using precast 10% TGX gels (Bio-Rad). Gels were run at 120 V and subsequently transferred to polyvinylidene fluoride (PVDF) membranes (BioRad) using BIO-RAD semi-dry transfer system. Then, membranes were fixed with 100% methanol for 1 minute and blocked with 5 % non-fat dry milk-containing Tris-buffered saline with Tween-20 (TBST) with 0.1% Tween-20 for 1 hour in shaking and room temperature (RT). Next, membranes were incubated with primary antibodies overnight at 4°C followed by incubation with secondary antibodies in a 3% milk diluted in TBST for 1 hour at RT. Primary antibodies against SARS-CoV-2 Spike S2 protein (Abcam; ab6823) and nucleocapsid (Novus Biologicals; NB100-56576) were purchased from the indicated suppliers and used at a dilution of 1:3000 and 1:2000 respectively. Anti-mouse secondary IgG-HRP antibody (abcam, 6823) was used at a dilution 1:5000 to detect SARS-CoV-2 Spike protein and anti-rabbit secondary IgG-HRP antibody (Kindle Biosciences, R1006) at 1:2000 to detect SARS-CoV-2 nucleocapsid.

### Plaque assay

To determine viral titers, 3.2 × 10^5^ VeroE6 or VeroE6-TMPRSS2 were seeded in a 12 well-plate the day before plaque assay was performed. Briefly, ten-fold serial dilutions were performed in infection media for SARS-CoV-2 and inoculated onto confluent VeroE6 or VeroE6-TMPRSS2 cell monolayer. After one-hour adsorption, supernatants were removed, and cells monolayers were overlaid with minimum essential media (MEM) containing 2% FBS and purified agar (OXOID) at a final concentration of 0.7%. Cells were then incubated 3 days at 37°C. Cells were fixed overnight with 10% formaldehyde for inactivation of potential SARS-CoV-2 virus. Overlay was removed and cells were washed once with PBS. Plaques were visualized by immunostaining. Briefly, cells were blocked in 5 % milk diluted in TBST. After 1-hour, anti-mouse SARS-CoV-2-NP antibody (1C7C7, kindly provided by Dr. Moran) was added at a dilution of 1:1000 in 1% milk-TBST and incubated for 1 hour at RT. Then, cells were washed two times in PBS and stained with goat anti-mouse secondary IgG-HRP antibody (abcam, 6823) at a dilution of 1:5000 in 1% milk-TBST and incubated for 1 hour at RT. Finally, cells were washed three times and the plaques were developed with TrueBlue substrate (KPL-Seracare). Viral titers were calculated as plaque forming units (PFU)/ml.

### RT-qPCR for viral infectivity analysis

.To quantify the levels of SARS-CoV-2 RNA after infection in VeroE6 and Caco-2 cells, we used the CDC 2019-nCoV real-time RT-qPCR protocol, with modifications. Primers and probes were purchased from the indicated supplier (Integrated DNA Technologies, 10006713, RUO Kit) and consisted of two 2019-nCoV-specific sets (N1, N2). Names and sequences of the primers used are shown in Supplementary Table 3. Assays were run in a 384-well format using the QuantiFast Pathogen RT-PCR + IC Kit (QIAGEN; 211454). USA/WA-1/2020 SARS-CoV-2 RNA (20,000 genome copies per reaction) and nuclease-free water were included as controls. Reactions were performed in duplicate using the following cycling conditions on the Roche LightCycler 480 Instrument II (Roche Molecular Systems; 05015243001): 50°C for 20 min, 95°C for 1 s, 95°C for 5 min, followed by 40 cycles of 95°C for 15 s and 60°C for 45 s. To determine the limit of detection for SARS-CoV-2, we used a commercially available plasmid control (Integrated DNA Technologies;10006625). Infectivity was calculated as a ratio between genomic RNA calculated by RT-qPCR and PFU values determined by plaque assay analysis.

### Immunofluorescence

VeroE6-TMPRSS2 cells were seeded at a concentration of 3.2 × 10^5^ cells per well in a 12 well glass-bottom plate and cultured at 37°C in 5% CO_2_ for 16 hours. Cells were then infected with the corresponding SARS-CoV-2 variant at an MOI of 0.01 and maintained in infection media. After 24 post-infection, cells were fixed with 10% methanol-free formaldehyde and incubated with primary antibodies against spike KL-S-3A7 (*43*) and nucleoprotein polyclonal anti-serum (*44*) diluted in 3% bovine serum albumin (BSA) for 1 hour at RT. Then, cells were washed and stained with secondary antibodies anti-mouse Alexa Fluor-488 (ThermoFisher; A21202) and anti-Rabbit Alexa Fluor 568 (ThermoFisher; A11011) in 5% BSA for 1h at RT. DAPI (4′,6-diamidino-2-phenylindole) was used to visualize the nucleus.

### Hamster infections

Ten female golden Syrian hamster of approximately 4-weeks-old were placed in pairs in five different cages. Only one hamster per cage was inoculated intranasally with a total of 10^5^ pfu of a mix of WA1 and WA1-655 SARS-CoV-2 viruses in a one-to-one ratio administered in 100 ul of PBS. Animals were monitored daily for body weight loss. On days 2, 4, 6 post-infection, animals were anesthetized with 100 mg/kg Ketamine and 20 mg/kg Xylazine and nasal washes were collected in 200 ul PBS. Directly infected (DI) and direct contact (DC) hamsters were humanely euthanized for collection of lungs and nasal turbinates on day 5 and 7 post-infection, respectively. Anesthetized hamsters were euthanized by intracardiac injection of sodium pentobarbital (Sleepaway - Zoetis) euthanasia solution. Samples were collected for viral quantification by plaque assay and next-generation sequencing. All animal studies were approved by the Institutional Animal Care and Use Committee (IACUC) of Icahn School of Medicine at Mount Sinai (ISMMS).

### Mink infection

Nine female American Mink (Neovison vison) of approximately 6-months-old were sourced by Triple F Farms (Gillett, PA). All mink were individually housed, given ad libitum access to food and water, and maintained on a 12-hour light/dark cycle. Six minks were infected with an infectious dose of 10^6^ pfu of WA-1 isolate administered intranasally in a 1 mL volume. Three minks were mock-infected to serve as healthy controls. Minks were anesthetized by intramuscular administration of 30 mg/kg Ketamine and 2 mg/kg Xylazine prior to intranasal infection, collection of specimens, or euthanasia. Nasal washes, rectal swabs, and oropharyngeal swabs were collected on days 1, 3, and 5 post-infection. On days 4 and 7 post-infection, three mink per day were humanely euthanized for collection of tissue specimens for viral quantification by plaque assay and sequencing. Body weights of mink were collected days 0, 1, 3, 4, 5 and 7 post-infection. Anesthetized minks were euthanized by intracardiac injection of sodium pentobarbital (Sleepaway - Zoetis) euthanasia solution. The Institutional Animal Care and Use Committee (IACUC) of the Icahn School of Medicine at Mount Sinai (ISMMS) reviewed and approved the mink model of COVID-19. Experiments with infected SARS-CoV-2 mink were performed in an ABSL-3 facility.

### Sample preparation for targeted proteomics

VeroE6-TMPRSS2 cells were seeded in a 6 well-plate and infected with the corresponding SARS-CoV-2 variants at an MOI of 0.1. Cells were lysated after 24 hours with RIPA buffer containing EDTA-free protease inhibitor cocktail and 10% SDS to a final concentration of 1%. Then, samples were boiled for 10 minutes at 100°C. Cells were centrifuged at 12k rpm on a tabletop centrifuge at RT for 20 minutes to remove insoluble debris and separated into three samples to assess technical reproducibility. 50 ul for each sample were loaded in a 1:4 ratio (v/v) with urea buffer (8M urea, 100 mM AmBic pH 8.1) on a Microcon 30 kDa MWCO (Millipore, Sigma) and centrifuged to dryness at 9500 rpm for 15 minutes at RT, until all sample was loaded. The filters were washed three times with 200 ul of urea buffer using similar centrifugation parameter as the sample loading. 100 ul of reduction buffer (8 M urea, 100 mM AmBic pH 8.1, 5 mM TCEP) was added and the samples were incubated at 37°C for 20 minutes to reduce the cysteines. Chloroacetamide (CAA) was added to 10 mM final concentration and the samples were incubated for 30 minutes in the dark at RT. The filters were washed 3 times with 200 ul of urea buffer and 3 times with 200 ul of digestion buffer (50 mM AmBiC). GluC was added to samples in a 1:100 ratio (w/w) and the filters were incubated on a shaker for 16 hours at 37°C and 450 rpm. Peptides were collected by centrifugation and the filters were washed twice with 100 ul of LC-MS grade water. Desalting was done using the Nest group microspin C18 columns. Activation of the resin was done with 1 column volume (CV) of MeCN and the columns were equilibrated with 2 CV of 0.1% FA in water. Samples were loaded and flowthrough was loaded again before washing the columns with 3 CV of 0.1% FA in water. Peptide elution was done with 2 CV of 50% MeCN in 0.1% FA and 1 CV of 80% MeCN in 0.1% FA. Following collection, the peptides were dried under the vacuum. Samples were resuspended at 1 ug/ul in 0.1% FA and approximately 1 ug was injected into the mass spectrometer.

### Mass spectrometry

All samples were acquired on a Thermo Q Exactive (Thermo Fisher) connected to a nanoLC easy 1200 (Thermo Fisher). Samples containing mutation in the region of interest (furin cleavage) as well as one representative sample having conservation in the furin cleavage site were analyzed by Data Dependent Acquisition (DDA) MS to obtain fragments library to design the targeted assays. For DDA the peptides were separated in 120 minutes with the following gradient: 4% B (0.1% FA in MeCN) to 18% B for 65 minutes, followed by another linear gradient from 18% to 34% of B and lastly the organic solvent was increased to 95% in 5 minutes and kept for 5 minutes to wash the column. The mass spectrometer was operated in positive mode, each MS1 scan was performed with a resolution of 70000 at 200 m/z. Peptide ions were accumulated for 100 ms or until the ion population reached an AGC of 3e6. The top 15 most abundant precursors were fragmented using high-collisional-induced dissociation (HCD) with a normalized energy of 27 using an isolation window of 2.2 Da and a resolution of 17500 (200 m/z). For targeted analysis, the samples were separated in 62 minutes to concentrate the analytes in narrower peaks and increase signal. The gradient employed was from 3% B to 34% in 40 minutes then B was increased to 42% in 10 minutes and then finally to 95% in 5 minutes. As for the DDA the column was washed for 5 minutes at 95% B before the next run. The mass spectrometer was operated in positive mode and targeted acquisition. Specifically, one MS1 scan (70000 resolution, 1e6 AGC, 100 ms IT) and seven unscheduled targeted MS2 scan were performed per cycle. Each MS2 was acquired at 35000 resolution, with a AGC of 2e5 a maximum IT of 110 ms and an isolation window of 2.0 m/z. Isolated ions were fragmented using HCD at 27 NCE.

### Illumina sequencing

Viral RNA nasal washes from hamsters was extracted using Omega E.Z.N.A Viral RNA kit (R6874) following manufacturer’s instructions. Viral RNA from hamster nasal turbinates and lungs was isolated Direct-zol RNA Miniprep kit (R2050) using manufacturer’s instructions. Samples for sequencing were prepared using whole-genome amplification with custom designed tiling primers (*45*) and the Artic Consortium protocol (https://artic.network/ncov-2019), with modifications. Briefly, cDNA synthesis was performed with random hexamers and ProtoScript II (New England Biolabs, cat. E6560) using 7 μL of RNA according to manufacturer’s recommendations. The RT reaction was incubated for 30 minutes at 48°C, followed by enzyme inactivation at 85°C for 5 minutes. Targeted amplification was performed as previously described(*31*). Next, amplicons were visualized on a 2% agarose gel and cleaned with Ampure XT beads. Amplicon libraries were prepared using the Nextera XT DNA Sample Preparation kit (Illumina, cat. FC-131-1096), as recommended by the manufacturer. Finally, to assembly SARS-CoV-2 genomes a custom reference-based analysis pipeline was used (https://github.com/mjsull/COVID_pipe ^43^). For the hamster samples and the inoculum, in addition to whole genome sequencing, the same workflow was used to sequence a 2,223 bp amplicon targeting the S1 to S1/S2 region (nucleotide positions 21,386 to 23,609) to quantify the variant frequency at the position S:655 position.

### Oxford Nanopore sequencing

The frequency of variants at position S:655 in the hamster samples was further confirmed with Oxford Nanopore (ONT) sequencing in a MinION Mk1C instrument. The same cDNA used for Illumina sequencing was used to amplify a fragment of 356bp along the spike region of interest (positions 23,468 to 23,821) using the Artic primer pair nCoV-2019_78 V3 (https://artic.network/ncov-2019). The purified amplicons were barcoded with the Native Barcode expansion kit (Oxford Nanopore, cat. EXP-NBD196) and PCR-free libraries were prepared using the ligation sequencing (Oxford Nanopore, cat. SQK-LSK109). A total of 20 fmol of the multiplexed library was sequenced on a Flongle flowcell (Oxford Nanopore, cat. FLO-FLG001) in a single 8-hour run. Sequencing data acquisition was done with the ONT software MinKNOW v4.3.7. Basecalling and demultiplexing was done with Guppy 5.0.12 in high accuracy basecalling mode. The Genome assembly was done with the Artic pipeline (artic-network/artic-ncov2019) with default parameters, where reads were aligned to the reference genome Wuhan-Hu-1 (MN908947.3) using minimap2 (2.17-r941), consensus variants were called with Nanopolish (0.13.2). Final read coverage for the targeted region ranged from 80 to 3251x, (median coverage of 1578x).

## Data Analysis

Multiple alignment of the spike protein of mink and human NY variants (Figure 1) was performed using COBALT multiple alignment tool (*46*). Spike amino acid sequence of SARS-CoV-2 ancestor (MN908947.3) was used to compare and identify the spike mutation of these variants. For the analysis of high prevalent amino acid changes within and surrounding the furin cleavage of the spike protein of current circulating VOCs, amino acid substitution frequencies around the cleavage site region (655 to 701) were estimated from globally available data (Available from GISAID [PMID: 31565258] as of 2021-06-28 with 2,072,987 records). The downloaded data was processed through the augur pipeline with Nextstrain v7 for SARS-CoV-2 (https://github.com/nextstrain/ncov) for sequence alignment and curation with default parameters (*47*). Multiple sequence alignment of the spike amino acid sequences was parsed and sliced for the region of interest in Biophyton v.75 with BioAlignIO package. In order to determine the more prevalent substitutions, sequences with ambiguous consensus calls were removed, and residues with substitutions along the 655-701 region present in frequencies of 0.05% or less across the entire dataset were masked. The 5% cutoff value was chosen based on the frequency distribution of substitutions per site on a histogram that revealed that variants in most positions occur in frequencies of less than 0.2%. The final list of most variable positions included residues 655, 675, 677, 681 and 701. The relative proportion of their occurrence time and Nextstrain clade (https://nextstrain.org/blog/2021-01-06-updated-SARS-CoV-2-clade-naming) was plotted in R with ggplot2 (*48*) v3.3.5.

Phylogenetic analysis was performed using the same dataset from above to build a time-calibrated phylogenetic tree with Nextstrain, to visualize the distribution of H655Y (up to 2020-09-30) and other prevalent substitutions along the 655-701 region (up to 2021-06-23). A global subsampling scheme with a focus on the variable residues was done to ensure their representation by geographical region and over time. The final builds contained 7,059 (early) and 13,847 (late) sequences including the early New York isolates.

For the mass spectrometry analysis DDA data was searched with MSFragger (*49*) using a FASTA file combining the human proteome, the SARS-Cov2 proteome, all the variants and the C-term and N-term cleaved Spike protein entry for each variant. The Speclib workflow was used to generate a library which was imported into Skyline (*50*) for selection of peptides and internal controls. Overall, 3 versions of the C-term furin cleaved peptide (SVASQSIIAYTMSLGAE) with two charge states (2+ and 3+) and the oxidated methionine were used. 4 other peptides were included as controls: 2 from the C-term spike fragment to be used as proxy for total spike quantity and 2 from Orf3a and N protein to be used as internal standard to normalize across variants. Following acquisition, the PRM data was imported into the Skyline document with the following transition settings: MS1 filtering was enabled, and MS/MS filtering was changed to targeted using Orbitrap as mass analyzer (35000 resolution) and high selectivity extraction. A minimum of 6 transitions and a maximum of 18 having m/z > precursors were selected for data analysis. After manual peak boundaries selection and elimination of interferences the transition results were exported. Transitions where the signal/background ratio was less than 5 were removed to ensure robust quantitative accuracy. The transitions were summed within the same charge state and the 2+ unmodified SVASQSIIAYTMSLGAE was used for quantification. The data was normalized using median centering of the other Spike peptide (ILPVSMTKTSVD) as internal standard. Following normalization, log_2_ fold change was calculated by averaging the intensities for the furin-cleaved peptide per variant and divide them by the one from the WA1 variant used here as control sample. Resulting ratios were logged and used for visualization and statistical analysis.

Statistical analysis was performed using GraphPad Prism. Two-way ANOVA with Holm-Šídák posttest was used for multiple comparisons. Statistical significance was established at a P value of <0.05.

## References

1. F. Fenollar et al., Mink, SARS-CoV-2, and the Human-Animal Interface. Front Microbiol 12, 663815 (2021).

2. J. Shi et al., Susceptibility of ferrets, cats, dogs, and other domesticated animals to SARS-coronavirus 2. Science 368, 1016–1020 (2020).

3. B. B. Oude Munnink et al., Transmission of SARS-CoV-2 on mink farms between humans and mink and back to humans. Science 371, 172–177 (2021).

4. A. S. Hammer et al., SARS-CoV-2 Transmission between Mink (Neovison vison) and Humans, Denmark. Emerg Infect Dis 27, 547–551 (2021).

5. P. Tong et al., Memory B cell repertoire for recognition of evolving SARS-CoV-2 spike. bioRxiv, (2021).

6. T. N. Starr et al., Deep Mutational Scanning of SARS-CoV-2 Receptor Binding Domain Reveals Constraints on Folding and ACE2 Binding. Cell 182, 1295–1310 e1220 (2020).

7. B. A. Johnson et al., Furin Cleavage Site Is Key to SARS-CoV-2 Pathogenesis. bioRxiv, (2020).

8. Y. Huang, C. Yang, X. F. Xu, W. Xu, S. W. Liu, Structural and functional properties of SARS-CoV-2 spike protein: potential antivirus drug development f or COVID-19. Acta Pharmacol Sin 41, 1141–1149 (2020).

9. L. Duan et al., The SARS-CoV-2 Spike Glycoprotein Biosynthesis, Structure, Function, and Antigenicity: Implications for the Design of Spike-Based Vaccine Immunogens. Front Immunol 11, 576622 (2020).

10. M. Hoffmann, H. Kleine-Weber, S. Pohlmann, A Multibasic Cleavage Site in the Spike Protein of SARS-CoV-2 Is Essential for Infection of Human Lung Cells. Mol Cell 78, 779–784 e775 (2020).

11. B. Coutard et al., The spike glycoprotein of the new coronavirus 2019-nCoV contains a furin-like cleavage site absent in CoV of the same clade. Antiviral Res 176, 104742 (2020).

12. D. Bestle et al., TMPRSS2 and furin are both essential for proteolytic activation of SARS-CoV-2 in human airway cells. Life Sci Alliance 3, (2020).

13. M. Ord, I. Faustova, M. Loog, The sequence at Spike S1/S2 site enables cleavage by furin and phospho-regulation in SARS-CoV2 but not in SARS-CoV1 or MERS-CoV. Sci Rep 10, 16944 (2020).

14. T. Tang et al., Proteolytic Activation of SARS-CoV-2 Spike at the S1/S2 Boundary: Potential Role of Proteases beyond Furin. ACS Infect Dis 7, 264–272 (2021).

15. S. Matsuyama et al., Efficient activation of the severe acute respiratory syndrome coronavirus spike protein by the transmembrane protease TMPRSS2. J Virol 84, 12658–12664 (2010).

16. S. Xia et al., Fusion mechanism of 2019-nCoV and fusion inhibitors targeting HR1 domain in spike protein. Cell Mol Immunol 17, 765–767 (2020).

17. J. A. Plante et al., Spike mutation D614G alters SARS-CoV-2 fitness. Nature 592, 116–121 (2021).

18. B. Korber et al., Tracking Changes in SARS-CoV-2 Spike: Evidence that D614G Increases Infectivity of the COVID-19 Virus. Cell 182, 812–827 e819 (2020).

19. Y. Liu et al., The N501Y spike substitution enhances SARS-CoV-2 transmission. bioRxiv, (2021).

20. C. Yi et al., Key residues of the receptor binding motif in the spike protein of SARS-CoV-2 that interact with ACE2 and neutralizing antibodies. Cell Mol Immunol 17, 621–630 (2020).

21. T. Aydillo et al., Immunological imprinting of the antibody response in COVID-19 patients. Nat Commun 12, 3781 (2021).

22. V. V. Edara et al., Infection and vaccine-induced neutralizing antibody responses to the SARS-CoV-2 B.1.617.1 variant. bioRxiv, (2021).

23. P. Mlcochova et al., SARS-CoV-2 B.1.617.2 Delta variant emergence, replication and sensitivity to neutralising antibodies. bioRxiv, 2021.2005.2008.443253 (2021).

24. D. Planas et al., Reduced sensitivity of SARS-CoV-2 variant Delta to antibody neutralization. Nature, (2021).

25. T. P. Peacock et al., The SARS-CoV-2 variants associated with infections in India, B.1.617, show enhanced spike cleavage by furin. bioRxiv, 2021.2005.2028.446163 (2021).

26. A. Baum et al., Antibody cocktail to SARS-CoV-2 spike protein prevents rapid mutational escape seen with individual antibodies. Science 369, 1014–1018 (2020).

27. K. M. Braun et al., Transmission of SARS-CoV-2 in domestic cats imposes a narrow bottleneck. PLoS Pathog 17, e1009373 (2021).

28. R. Rathnasinghe et al., The N501Y mutation in SARS-CoV-2 spike leads to morbidity in obese and aged mice and is neutralized by convalescent and post-vaccination human sera. medRxiv, (2021).

29. Z. Liu et al., Identification of Common Deletions in the Spike Protein of Severe Ac ute Respiratory Syndrome Coronavirus 2. J Virol 94, (2020).

30. M. M. Lamers et al., Human airway cells prevent SARS-CoV-2 multibasic cleavage site cell culture adaptation. Elife 10, (2021).

31. A. S. Gonzalez-Reiche et al., Introductions and early spread of SARS-CoV-2 in the New York City area. Science 369, 297–301 (2020).

32. N. N. Gaudreault et al., SARS-CoV-2 infection, disease and transmission in domestic cats. Emerg Microbes Infect 9, 2322–2332 (2020).

33. S. F. Sia et al., Pathogenesis and transmission of SARS-CoV-2 in golden hamsters. Nature 583, 834–838 (2020).

34. K. Rosenke et al., Defining the Syrian hamster as a highly susceptible preclinical model for SARS-CoV-2 infection. Emerg Microbes Infect 9, 2673–2684 (2020).

35. W. T. Harvey et al., SARS-CoV-2 variants, spike mutations and immune escape. Nat Rev Microbiol 19, 409–424 (2021).

36. N. Rego et al., Spatiotemporal dissemination pattern of SARS-CoV-2 B1.1.28-derived lineages introduced into Uruguay across its southeastern border with Brazil. medRxiv, 2021.2007.2005.21259760 (2021).

37. S. Banerjee, S. Seal, R. Dey, K. K. Mondal, P. Bhattacharjee, Mutational spectra of SARS - CoV-2 orf1ab polyprotein and signature mutations in the United States of America. J Med Virol 93, 1428–1435 (2021).

38. F. Begum et al., Specific mutations in SARS-CoV2 RNA dependent RNA polymerase and helicase alter protein structure, dynamics and thus function: Effect on viral RNA replication. (2020).

39. J. Buchrieser et al., Syncytia formation by SARS-CoV-2-infected cells. EMBO J 39, e106267 (2020).

40. X. Chi et al., A neutralizing human antibody binds to the N-terminal domain of the Spike protein of SARS-CoV-2. Science 369, 650–655 (2020).

41. J. Lan et al., Structure of the SARS-CoV-2 spike receptor-binding domain bound to the ACE2 receptor. Nature 581, 215–220 (2020).

42. T. Aydillo et al., Shedding of Viable SARS-CoV-2 after Immunosuppressive Therapy for Cancer. N Engl J Med 383, 2586–2588 (2020).

43. F. Amanat et al., Murine Monoclonal Antibodies against the Receptor Binding Domain of SARS-CoV-2 Neutralize Authentic Wild-Type SARS-CoV-2 as Well as B.1.1.7 and B.1.351 Viruses and Protect In Vivo in a Mouse Model in a Neutralization-Dependent Manner. mBio, e0100221 (2021).

44. M. Spiegel et al., Inhibition of Beta interferon induction by severe acute respiratory syndrome coronavirus suggests a two-step model for activation of interferon regulatory factor 3. J Virol 79, 2079–2086 (2005).

45. J. Quick et al., Multiplex PCR method for MinION and Illumina sequencing of Zika and other virus genomes directly from clinical samples. Nat Protoc 12, 1261–1276 (2017).

46. J. S. Papadopoulos, R. Agarwala, COBALT: constraint-based alignment tool for multiple protein sequences. Bioinformatics 23, 1073–1079 (2007).

47. J. Hadfield et al., Nextstrain: real-time tracking of pathogen evolution. Bioinformatics 34, 4121–4123 (2018).

48. H. Wickham, ggplot2: Elegant Graphics for Data Analysis. https://ggplot2.tidyverse.org. Springer, New York; 2016.

49. A. T. Kong, F. V. Leprevost, D. M. Avtonomov, D. Mellacheruvu, A.I. Nesvizhskii, MSFragger: ultrafast and comprehensive peptide identification in mass spectrometry-based proteomics. Nat Methods 14, 513–520 (2017).

50. B. MacLean et al., Skyline: an open source document editor for creating and analyzing targeted proteomics experiments. Bioinformatics 26, 966–968 (2010).

