## Supplementary Materials for "SARS-CoV-2 variants of concern have acquired mutations associated with an increased spike cleavage"

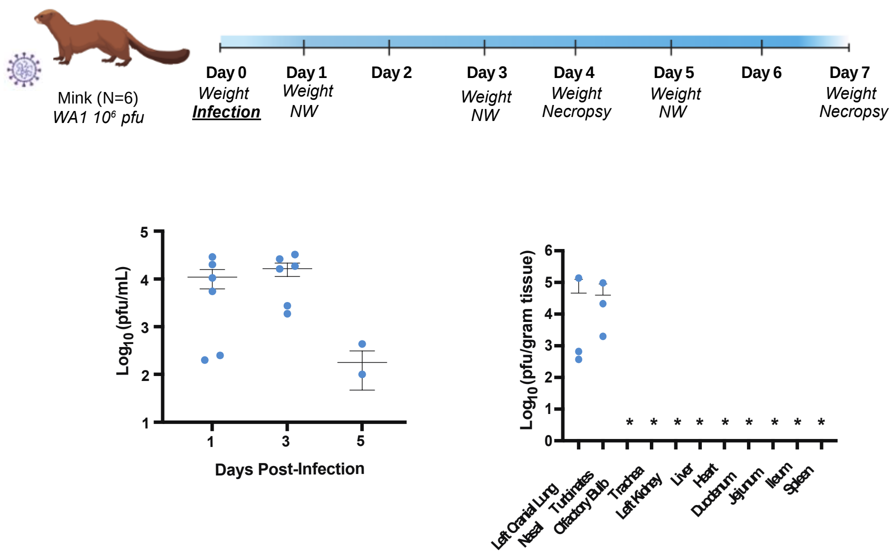


**Fig. S1. Spike polymorphism H655Y is selected after SARS-CoV-2 replication in mink in vivo model.** A) Mink study design: six minks were infected with 106 pfu of SARS-CoV-2 WA1 isolate. Nasal washes were collected at day 1, 3 and 5 post-infection (p.i.) and organs were harvested at day 3 and 7 p.i. B) Viral titers of nasal washes expressed as PFU per milliliter. C) Organ viral titers expressed as pfu per gram of tissue.

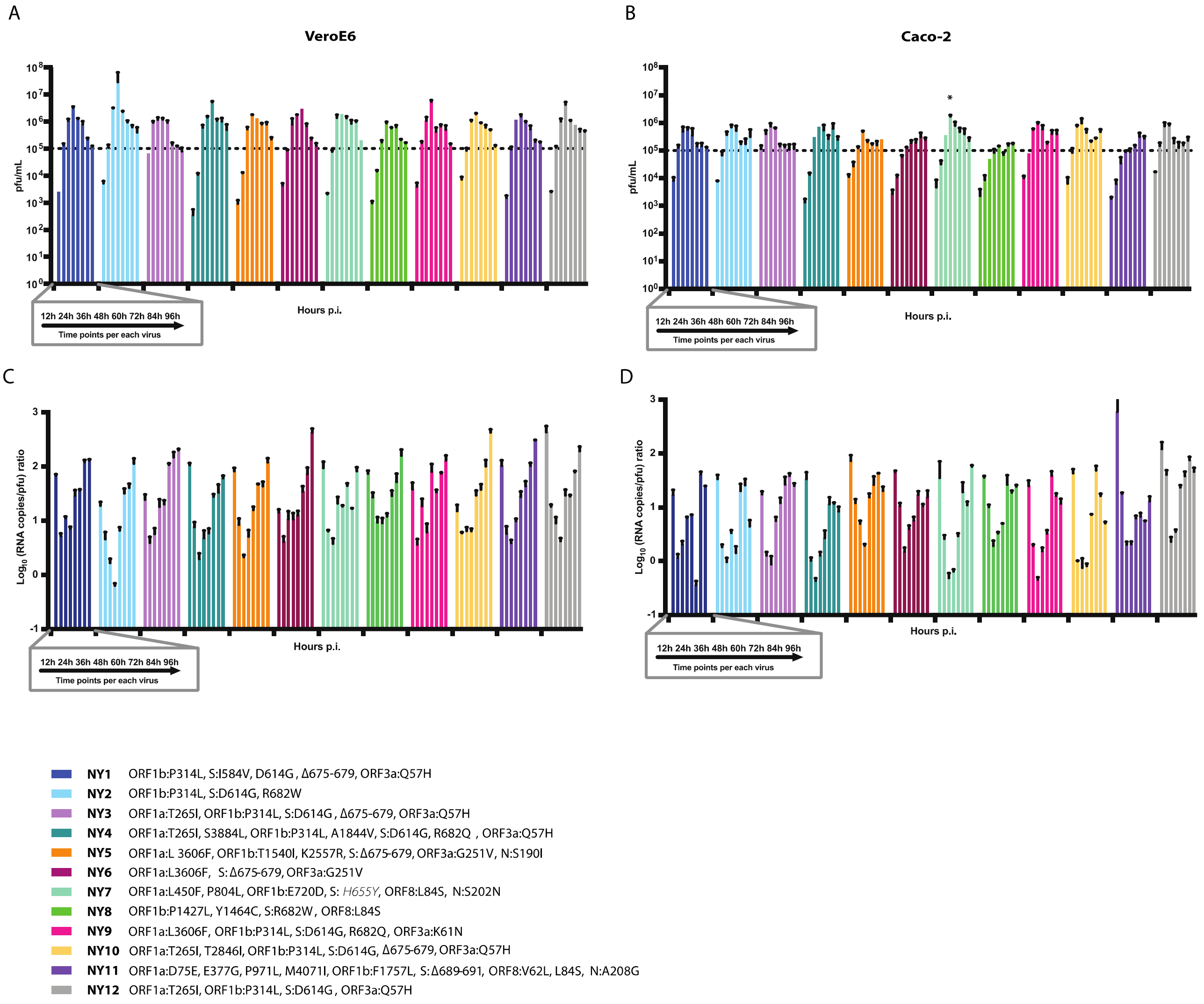


**Fig. S2. Spike polymorphism H655Y identified in SARS-CoV-2 NY7 confers a growth advantage in human Caco-2 cells.** Replication kinetics of 12 NY human isolates in VeroE6 A) and Caco-2 cells B). Cells were infected at a multiplicity of infection (MOI) of 0.01. Viral titers were determined by plaque assay and expressed as PFU per milliliter at the indicated hours post-infection (p.i.). Means and error bars are shown. C). Virion infectivity expressed as genomic RNA/pfu ratio in VeroE6 C) and Caco-2 cells D). Cells were infected at a MOI of 0.01. The genomic viral RNA was determined by RT-qPCR. Shown are the means and error bars represent standard deviations.

**Table S1.** GISAID accession numbers of human SARS-CoV-2 variants isolated from New York (NY) during the first COVID-19 pandemic outbreak.

| SARS-CoV-2 isolate | GISAID accession number |
|---|---|
| NY1 | HCOV19/NY/PV08139/2020_EPI_ISL_MSPV08139 |
| NY2 | HCOV19/NY/PV08426/2020_EPI_ISL_MSPV08426 |
| NY3 | HCOV19/NY/PV08428/2020_EPI_ISL_MSPV08428 |
| NY4 | HCOV19/NY/PV08485/2020_EPI_ISL_MSPV08485 |
| NY5 | HCOV19/NY/PV08137/2020_EPI_ISL_MSPV08137 |
| NY6 | HCOV19/NY/PV08489/2020_EPI_ISL_MSPV08489 |
| NY7 | HCOV19/NY/PV08148/2020_EPI_ISL_MSPV08148 |
| NY8 | HCOV19/NY/PV08146/2020_EPI_ISL_MSPV08146 |
| NY9 | HCOV19/NY/PV08462/2020_EPI_ISL_MSPV08462 |
| NY10 | HCOV19/NY/PV08468/2020_EPI_ISL_MSPV08468 |
| NY11 | HCOV19/NY/PV08490/2020_EPI_ISL_MSPV08490 |
| NY12 | HCOV19/NY/PV08495/2020_EPI_ISL_MSPV08495 |

**Table S2.** Genomic mutations of mink-derived SARS-CoV-2 isolates (MiA) and human SARS-CoV-2 circulating variants during the first pandemic outbreak in New York (NY).

| SARS-CoV-2 isolate | Mutations |
|---|---|
| MiA-1 | ORF1a:S692P, S:T259K, H655Y, H1159Y, ORF8:L84S |
| MiA-2 | S:T259K, H655Y, H1159Y, ORF8:L84S |
| MiA-3 | S:T259K, H655Y, ORF6:F22*, ΔK23, ORF8:L84S |
| MiA-4 | S:H655Y, R682W, ORF8:L84S |
| MiA-5 | ORF1a:L3606F, S:H655Y, R682W, ORF8:L84S |
| MiA-6 | S:H655Y, R682W, ORF8:L84S |
| NY1 | ORF1b:P314L, S:I584V, D614G, Δ675-679, ORF3a:Q57H |
| NY2 | ORF1b:P314L, S:D614G, R682W |
| NY3 | ORF1a:T265I, ORF1b:P314L, S:D614G, Δ675-679, ORF3a:Q57H |
| NY4 | ORF1a:T265I, S3884L, ORF1b:P314L, A1844V, S:D614G, R682Q, ORF3a:Q57H |
| NY5 | ORF1a:L3606F, ORF1b:T1540I, K2557R, S:Δ675-679, ORF3a:G251V, N:S190I |
| NY6 | ORF1a:L3606F, S:Δ675-679, ORF3a:G251V |
| NY7 | ORF1a:L450F, P804L, ORF1b:E720D, S:H655Y, ORF8:L84S, N:S202N |
| NY8 | ORF1b:P1427L, Y1464C, S:R682W, ORF8:L84S |
| NY9 | ORF1a:L3606F, ORF1b:P314L, S:D614G, R682Q, ORF3a:K61N |
| NY10 | ORF1a:T265I, T2846I, ORF1b:P314L, S:D614G, Δ675-679, ORF3a:Q57H |
| NY11 | ORF1a:D75E, E377G, P971L, M4071I, ORF1b:F1757L, S:Δ689-691, ORF8:V62L, L84S, N:A208G |
| NY12 | ORF1a:T265I, ORF1b:P314L, S:D614G, ORF3a:Q57H |

**Table S3.** Primers used to detect SARS-CoV-2 RNA in nasopharyngeal swabs from COVID-19 infected patients.

| **Primer** | **Nucleotide sequence 5’>3’** |
|---|---|
| 2019-nCoV_N1 Forward Primer | GAC CCC AAA ATC AGC GAA AT |
| 2019-nCoV_N1 Reverse Primer | TCT GGT TAC TGC CAG TTG AAT CTG |
| 2019-nCoV_N2 Forward Primer | TTA CAA ACA TTG GCC GCA AA |
| 2019-nCoV_N2 Reverse Primer | GCG CGA CAT TCC GAA GAA |
